# Acidic nanoparticles restore lysosomal acidification and rescue metabolic dysfunction in pancreatic β-cells under lipotoxic condition

**DOI:** 10.1101/2023.07.11.548395

**Authors:** Chih Hung Lo, Lance M. O’Connor, Gavin Wen Zhao Loi, Eka Norfaishanty Saipuljumri, Jonathan Indajang, Kaitlynn M. Lopes, Orian S. Shirihai, Mark W. Grinstaff, Jialiu Zeng

**Author notes:** Correspondence: Jialiu Zeng, PhD Chih Hung Lo, PhD. Equal contribution (Chih Hung Lo, Jialiu Zeng).

## Abstract

Type 2 diabetes (T2D), a prevalent metabolic disorder lacking effective treatments, is associated with lysosomal acidification dysfunction as well as autophagic and mitochondrial impairments. Here, we report a series of biodegradable poly(butylene tetrafluorosuccinate-co-succinate) (PBFSU) polyesters, comprising an 1,4-butanediol linker and varying ratios of tetrafluorosuccinic acid (TFSA) and succinic acid as components, to engineer new lysosome acidifying nanoparticles (NPs). Notably, TFSA NPs, which composed entirely of TFSA, exhibit the strongest degradation capability and superior acidifying property. We further reveal significant downregulation of lysosomal vacuolar (H+)-ATPase (V-ATPase) subunits, which are responsible for maintaining lysosomal acidification, in human T2D pancreatic islets and INS-1 β-cells under lipotoxic condition. Treatment of TFSA NPs counteracts lipotoxicity in INS-1 β-cells by restoring lysosomal acidification, autophagic function, and mitochondrial activity, along with promoting glucose-stimulated insulin secretion. Administration of TFSA NPs to high-fat diet T2D mice improves glucose clearance and reduces insulin resistance. These findings highlight the therapeutic potential of lysosome acidifying TFSA NPs for T2D.

**Graphical Table of Contents:** 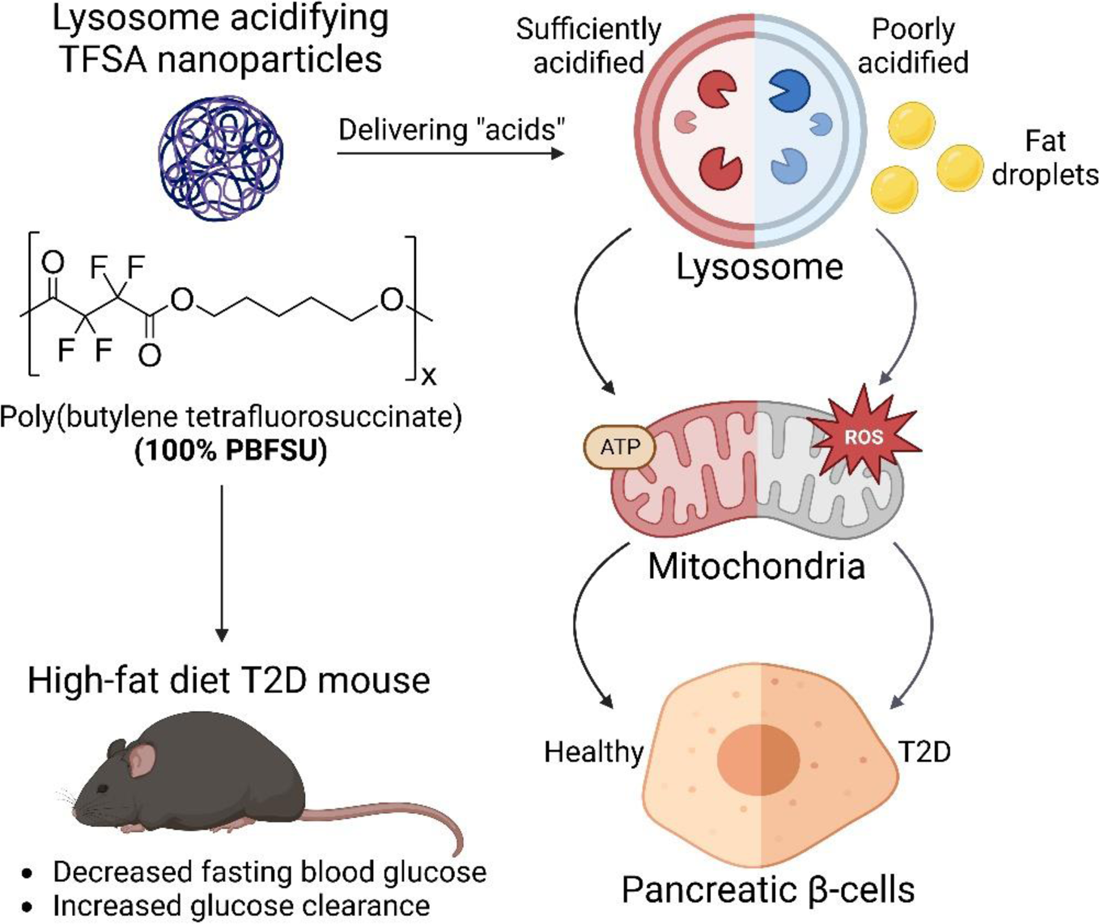

## Introduction

Diabetes is a metabolic disease characterized by an elevated body mass index and high levels of fasting blood glucose (1). Long-term complications are significant in diabetics and include heart attack, kidney failure, and blindness, among others (1). Currently, approximately 537 million adults live with diabetes, and the total number of diabetic patients is projected to rise to 643 million by 2030 and 783 million by 2045 (2). Type 2 diabetes (T2D) is the most common type of diabetes, afflicting 90-95% of patients (3). Obesity-related T2D generally associates with reduced mitochondrial activities and insulin secretory function in pancreatic β-cells as well as insulin resistance in other cell types such as hepatocytes, resulting in elevated fasting plasma insulin and blood glucose levels (4, 5).

Current clinical treatments for T2D include metformin, which lowers blood glucose levels via inhibition of liver gluconeogenesis and increases insulin sensitivity through enhanced glucose uptake in the skeletal muscles and in the gut (6, 7). However, due to its pleiotropic effects, metformin results in severe side effects, including lactic acidosis (8). Other class of drugs includes sulfonylureas, which close the adenosine triphosphate (ATP)-sensitive potassium channels in the cell membrane of the pancreatic β-cells and induces membrane depolarization, subsequently leading to calcium influx and insulin release (9). However, sulfonylureas treatment also induces side effects such as hypoglycemia and is only effective with intact functional pancreatic β-cells (10). Incretin mimetics such as exenatide and liraglutide have also been used to stimulate the release of insulin in response to glucose (4, 11). For instance, exenatide is an agonist which binds to and activate glucagon-like peptide-1 (GLP-1) receptors on pancreatic β-cells to increase insulin secretion (12, 13). However, the GLP-1 receptor agonist suffers from short plasma half-lives which leads to multiple dosing, thereby increasing the cost of treatment (13). Hence, there is an imperative need to understand the underlying disease mechanism related to T2D, as well as to develop more effective therapeutics.

Lysosomal function and its impairment are closely tied to obesity-induced T2D. Chronic accumulation of free fatty acids, such as palmitic acid, in pancreatic β-cells impairs lysosomal acidification and function, thereby causing autophagic and mitochondrial dysfunction (**Figure 1A**) (14–17). Thus, targeting lysosomal function to promote autophagy and mitophagy activities is a promising approach for treatment of T2D (18, 19). GLP-1 receptor agonist treatment increases transcription factor EB (TFEB) nuclear translocation, thereby increasing lysosomal biogenesis and lysosomal function and promoting β-cells survival under lipotoxic condition (15). Our recent work shows that treatment with lysosome-targeting nanoparticles (NPs), such as photo-activated NPs (PaNPs) (16, 20) and poly(lactic-co-glycolic acid) (PLGA) NPs (17), directly lowers lysosome luminal pH and restores lysosomal acidification and function, leading to enhanced autophagic function and mitochondrial function as well as increased insulin secretion from β-cells under lipotoxic condition. However, PaNPs require the application of an ultraviolet light trigger and PLGA NPs require a high concentration (i.e., >1 mg/mL) to be effective as they are weakly acidic (17, 21, 22). Both NP formulations are non-ideal for *in vivo* use and document the need for more effective lysosome acidifying therapeutics.

**Figure 1.**
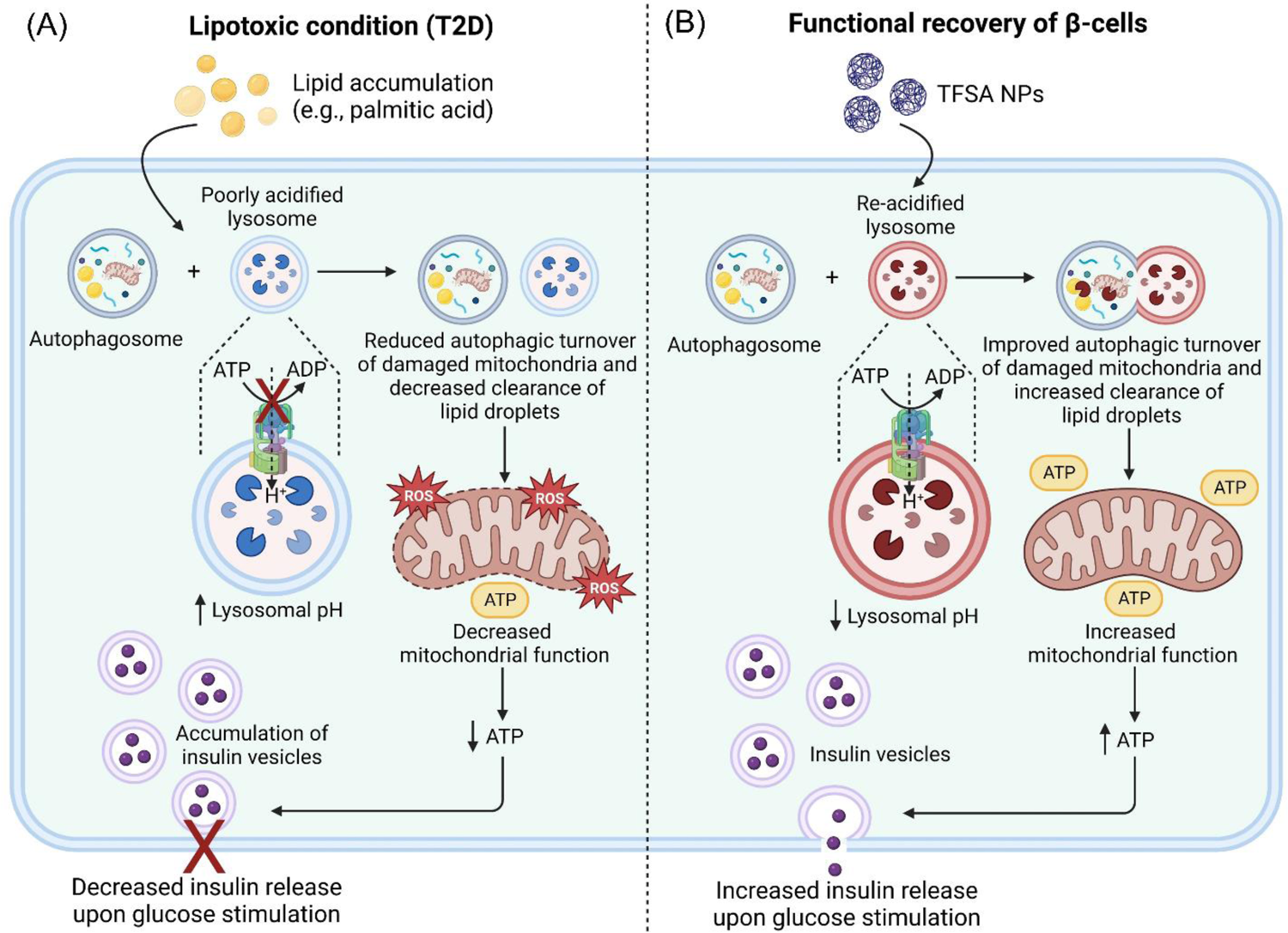
Schematic representation of fatty acid induced lipotoxicity in β-cells and functional recovery of lysosomal and cellular functions with treatment of TFSA NPs. (A) Under lipotoxic condition or lipid accumulation such as palmitic acid treatment, the lysosomal pH is elevated, and lysosomal functions are impaired. This causes a reduction in autophagic turnover of damaged mitochondria and a decrease in clearance of lipid droplets. The accumulation of impaired mitochondria which have decreased function leads to a lower ATP level, resulting in intracellular accumulation of insulin vesicles and a decrease in insulin release upon glucose stimulation. (B) Treatment of TFSA NPs re-acidify impaired lysosomes and promote their functions in β-cells under lipotoxic condition which leads to improved autophagic turnover of damaged mitochondria and increased clearance of lipid droplets. This results in increased mitochondrial function that produces more ATP and increases insulin release upon glucose stimulation.

In this study, we describe a new series of biodegradable poly(butylene tetrafluorosuccinate-co-succinate) (PBFSU) polyesters composed of 1,4-butanediol and varying ratios of tetrafluorosuccinic acid (TFSA) and succinic acid (SA) as components to engineer new lysosome acidifying NPs. Specifically, the NPs generated from the polyesters made up of entirely 100% TFSA (TFSA NPs) exhibit the high degree of component acid release to impart the strongest effect in lysosomal acidification. In fact, downregulation of lysosomal vacuolar (H^+^)-ATPase (V-ATPase) subunits in mRNA expression of human T2D islets from deposited dataset and in protein expression of insulinoma INS-1 pancreatic β-cells under palmitate induced lipotoxic condition confirm defective lysosomal acidification in obesity-induced T2D. TFSA NPs restore lysosomal acidification impairment, improve autophagic and mitochondrial functions, and increase insulin secretion under glucose stimulation in INS-1 β-cells (**Figure 1B**). In mice under high-fat diet (HFD), treatment with TFSA NPs reduces plasma insulin and glucose levels as well as improves glucose clearance, demonstrating a decrease in insulin resistance and more efficient utilization of insulin. TFSA NPs represent a new class of lysosome acidifying therapeutics for T2D and obesity induced metabolic diseases in general.

## Results and Discussion

### Engineering of TFSA NPs with lysosome acidifying property

To engineer NP-based lysosome acidifying agents, we designed polyesters that efficiently form into NPs and are capable of degrading in mildly acidic aqueous conditions of pH 5.5 to 6 to release the component acids. TFSA with a pKa of ∼1.6 and SA with a pKa of ∼4.2 are used as component acids (23) and the percentage of each acid used in synthesizing the polyesters controls the acidification capability of the subsequently formed NPs. In addition, fluorinated polyesters, in general, exhibit *in vitro* and *in vivo* biocompatibility (24), and are used in medical imaging and drug delivery devices. Furthermore, the linker used in the synthesis of the polyesters, 1,4-butanediol, is a common excipient used in the pharmaceutical industry and a component of dietary supplements (25).

We varied the ratios of TFSA to SA to synthesize a range of PBFSU polyesters from 0% PBFSU (also known as poly(butylene succinate) (PBSU)) to 25%-100% PBFSU using a dehydration polycondensation reaction (**Figure 2A**). The compositions of all synthesized polyesters were determined by nuclear magnetic resonance (NMR) spectroscopy (**Figure S1, S2, and S3**) and their molecular weights were measured by gel permeation chromatography (GPC) (**Figure S4A**). We also determined the melting temperature (T_m_), crystallization temperature (T_c_), glass transition temperature (T_g_), and the temperature at which the polyesters show 5% degradation (T_d,5%_) or maximum degradation (T_d,max_) of all synthesized polyesters by differential scanning calorimetry (DSC) and thermogravimetric analysis (TGA) (**Figure S4B**). Based on their thermal properties, the PBSFU polyesters are classified into two types according to their melt crystallizability where PBSU and 25% PBFSU polyesters are crystalline and 50-100% PBFSU polyesters are amorphous. All the polyesters exhibit a single glass transition temperature, which indicates that they are random copolymers (26).

**Figure 2.**
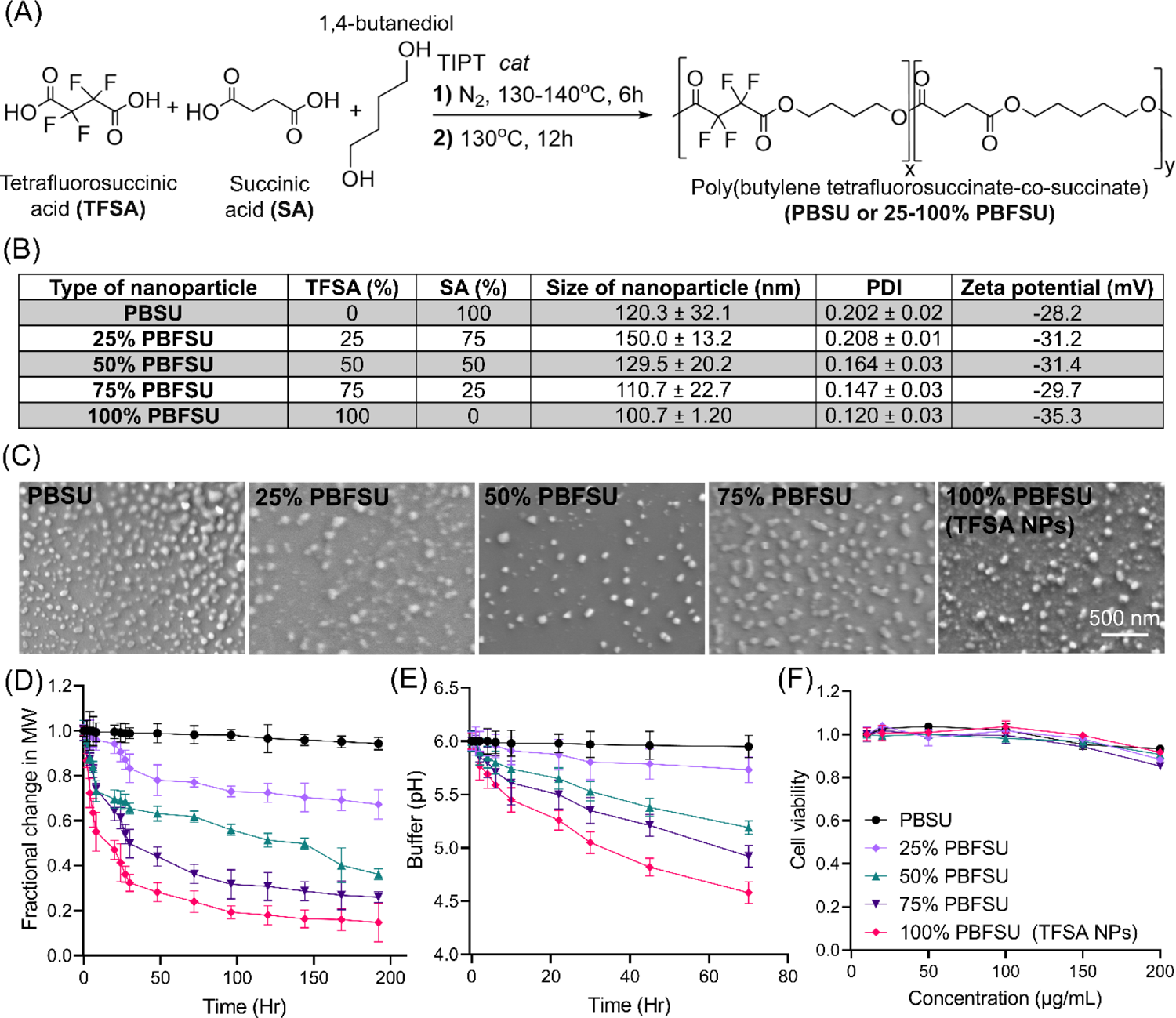
Engineering of TFSA nanoparticles (NPs) consisting of 100% PBFSU polyester that have strong degradation capability and superior acidifying property. (A) Scheme for the synthesis of poly (butylene tetrafluorosuccinate-co-succinate) (PBSU or 25-100% PBFSU) polyesters from tetrafluorosuccinic acid (TFSA) and succinic acid (SA) as well as an 1,4-butanediol linker. Various types of NPs were engineered based on different component acids through nanoprecipitation method. (B) Characterization of NPs size, polydispersity (PDI), and zeta potential. (C) Scanning electron microscopy (SEM) images to characterize the shape and morphology of the different types of NPs. NPs consisting of 100% PBFSU were termed TFSA NPs. (D) Gel permeation chromatography (GPC) analysis to determine the degradation rate of the different types of NPs with TFSA NPs (red line) showing the strongest degradation capability. (E) Determination of the extent of buffer acidification by the different types of NPs with TFSA NPs (red line) demonstrating superior acidifying property under mildly acidic condition of pH 6.0. (F) Cell cytotoxicity assay illustrating that all types of NPs, including TFSA NPs (red line), are non-toxic to INS-1 β-cells. Data are means ± SD of N=3 independent experiments.

Amongst the different NP fabrication techniques such as mini-emulsion (27), solvent displacement (28), and nanoprecipitation (29, 30), we selected nanoprecipitation as it is a facile and efficient method that does not require high shear stress for NP preparation, and is easily scalable (29, 30). Moreover, NPs formed through the nanoprecipitation method have also been reported to adopt a smaller size compared to other fabrication methods (29, 30). As the surfactant concentration relative to polymer concentration directly influences NP size and stability (31, 32), we varied the polymer to surfactant concentration using 100% PBFSU polyester and determined the optimal ratio to be 1:4 for the formation of NPs that are around 100 nm with a low polydispersity (**Figure S5A-B**). We then engineered the full series of PBFSU based NPs with them being spherical, having low polydispersity, and illustrating high zeta potential which indicates high stability (**Figure 2B-C**). We then determined the degradation capability (**Figure 2D**) and acidification strength (**Figure 2E and S5C**) of all PBFSU based NPs. Importantly, the 100% PBFSU, which we termed TFSA NPs, possess the highest degree of degradation into the component acids (**Figure 2D**) and the strongest buffer acidification strength at mildly acidic environment of pH 6.0 (**Figure 2E**) but not at neutral condition of pH 7.4 (**Figure S5C**). Finally, we confirmed that PBFSU based NPs, including TFSA NPs, are non-toxic to INS-1 β-cells (**Figure 2F**), and thus suitable to be tested in INS-1 β-cells for their effect in restoring lysosomal acidification.

### Presence of lysosomal dysfunction in human T2D islets

To investigate lysosomal, autophagic, and mitochondrial dysfunction associated with T2D in human, we conducted data mining to analyze an existing microarray transcriptome dataset (GSE25724) containing mRNA expression of T2D (6 disease samples) and non-diabetic (7 control samples) isolated human islets (33). With a cutoff of adjusted *P*<0.05 and log_2_(fold change (FC))>|1|, we identified 1448 downregulated and 78 upregulated differentially expressed genes (DEGs) as indicated in the volcano plot (**Figure 3A**) and in the heatmap (**Figure 3B**) from a total of 22283 genes. We then performed pathway enrichment analysis to obtain the functional annotations of the DEGs. From the pathway enrichment analysis using Kyoto Encyclopedia of Genes and Genomes (KEGG), the identified DEGs are associated with changes in metabolic pathways, tricarboxylic acid (TCA) cycle, proteasome and ubiquitin mediated proteolysis, protein processing in endoplasmic reticulum (ER), as well as endocytosis, autophagy, and lysosomal functions (**Figure 3C**). The DEGs are also related to diabetic cardiomyopathy, Alzheimer’s disease, Parkinson’s disease, and pathways of neurodegeneration, which are the diseases that have been shown to illustrate metabolic defects and interplay with one another (**Figure 3C**) (34, 35). From the pathway enrichment analysis using gene ontology biological process (GOBP), we additionally show alterations in biological processes that are associated with fatty acid and amino acid metabolism, catabolic processes, protein transport, and insulin processing (**Figure 3D**). Importantly, DEGs are highly enriched in autophagic and lysosomal functions, especially related to regulation of macroautophagy and vacuolar acidification (**Figure 3D**). We then identified seven lysosomal V-ATPase subunits (*ATP6V1B2*, *ATP6AP2*, *ATP6V1A*, *ATP6V1H*, *ATP6V1G1*, *ATP6V0E1*, and *ATP6V0B*) to be significantly downregulated in the dataset (**Figure 3E**), confirming that lysosomal functions are impaired in human T2D islets.

**Figure 3.**
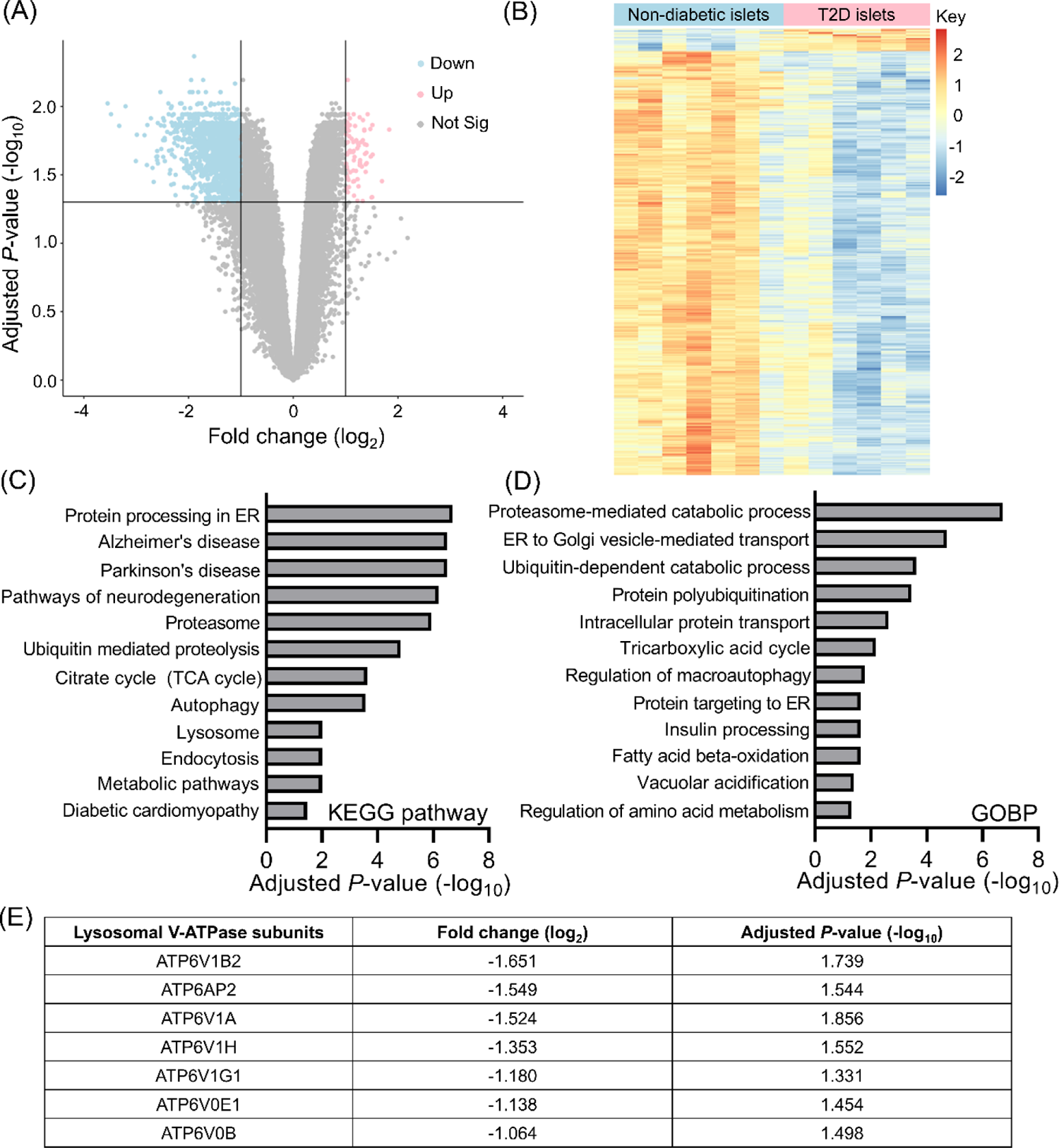
Data mining of microarray transcriptome dataset from GSE25724 containing mRNA expression of human T2D and non-diabetic islets. (A) Volcano plot illustrating the differentially expressed genes (DEGs) from the GSE25724 dataset based on the mRNA expression of the T2D (6 samples) and non-diabetic (7 samples) islets with a cut-off of adjusted *P*<0.05 and log_2_(fold change)>|1|. (B) Heatmap illustrating the mRNA profiles of T2D (6 samples) and non-diabetic (7 samples) islets from the GSE25724 dataset. (C) Kyoto Encyclopedia of Genes and Genomes (KEGG) analysis showing the enriched pathways associated with the DEGs. (D) Gene ontology biological process (GOBP) analysis showing the enriched biological processes associated with the DEGs. (E) Specific lysosomal V-ATPase subunits that are identified to be downregulated in T2D islets. All analyses were performed with adjusted *P*< 0.05.

### TFSA NPs restore lysosomal acidification and autophagic function in a palmitate-treated INS-1 cellular model of lipotoxicity

While fatty acid, specifically palmitic acid, treated INS-1 β-cells are widely used as a cellular model for T2D, the role of lysosomal dysfunction in these cells under lipotoxic condition remains unclear. To clarify this, we probed for the top three downregulated (log_2_FC<-1.5) lysosomal V-ATPase subunits (ATP6V1B2, ATP6AP2, and ATP6V1A) in palmitate treated INS-1 β-cells and observed a significant reduction in the protein expression of these subunits (**Figure 4A-B**), recapitulating their downregulation in human T2D islets (**Figure 3E**) and making this cell model suitable to study lysosomal acidification related dysfunction. Palmitate induces lysosomal pH elevation from 4.3 to 5.4 in INS-1 β-cells as compared to the fatty acid-free bovine serum albumin (BSA) control, and treatment of TFSA NPs acidifies the impaired lysosomes by lowering their pH from 5.4 to 4.5 (**Figure 4C-D**). Palmitate exposure also impairs lysosomal cathepsin L activity which requires a sufficiently acidic lysosome for optimal functioning, and TFSA NPs treatment restores enzyme activity (**Figure 4E**). Further, palmitate inhibits the autophagic function as shown by the accumulation of p62 and LC3II in INS-1 β-cells (36), which is reversed with TFSA NPs treatment (**Figure 4F-H**).

**Figure 4.**
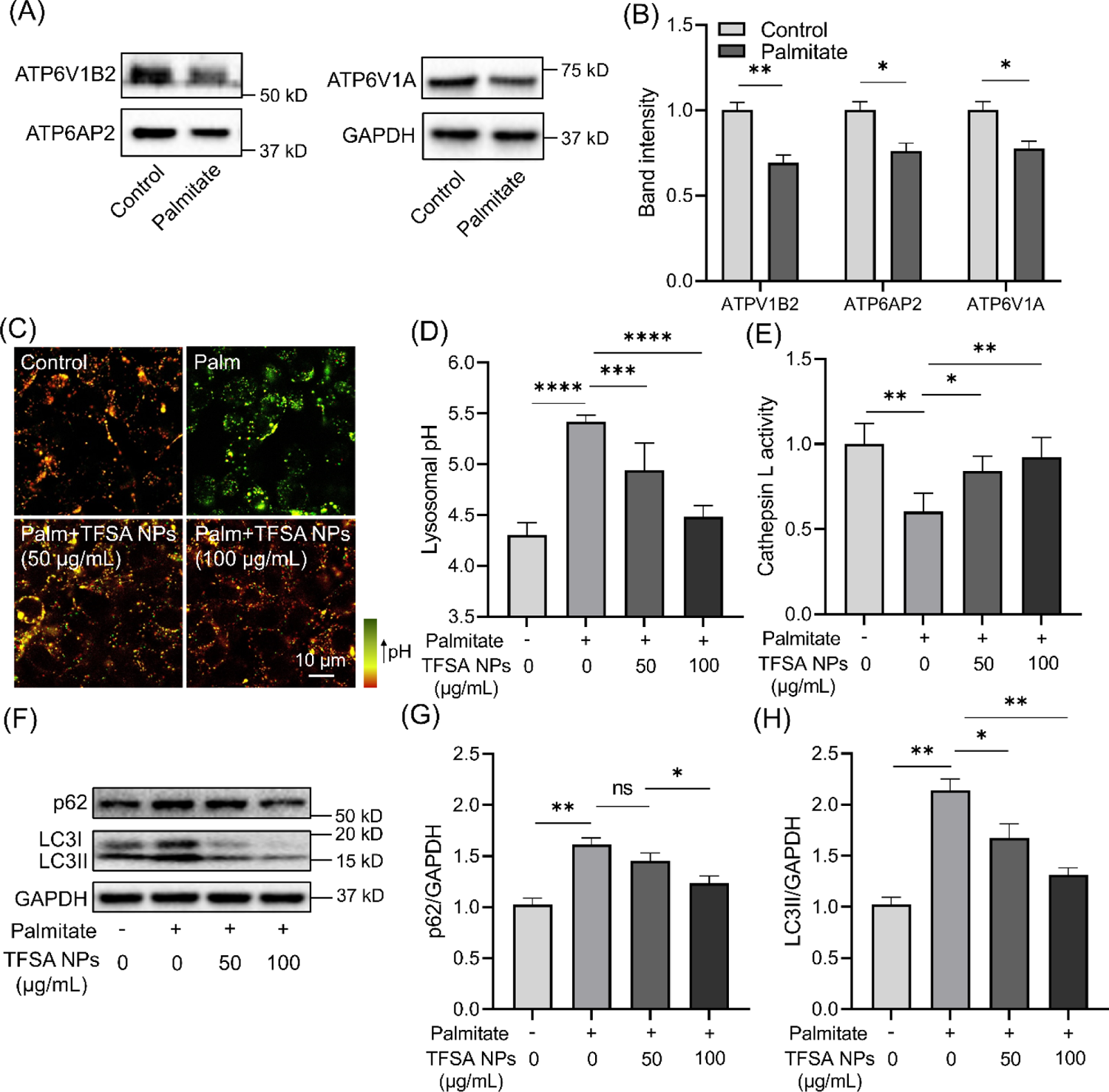
TFSA NPs restore lysosomal acidification and autophagic function in INS-1 β-cells under lipotoxic condition with palmitate treatment. (A-B) Western blotting characterization and quantification showing reduced lysosomal V-ATPase subunits in INS-1 β-cells under lipotoxic condition with palmitate treatment. (C-D) Immunofluorescence and quantification illustrating an elevated lysosomal pH in palmitate treated INS-1 β-cells and addition of TFSA NPs lowers lysosomal pH and restores luminal acidification in these cells. (E) TFSA NPs restore lysosomal cathepsin L function in palmitate treated INS-1 β-cells. (F-H) Western blotting characterization and quantification demonstrating autophagic function impairment as characterized by the accumulation of p62 (G) and LC3II (H) in palmitate treated INS-1 β-cells, and TFSA NPs promote autophagic degradation under lipotoxic condition. Data presented are relative to control without palmitate and TFSA NPs. Data are means ± SD of N=4 independent experiments. \**P* < 0.05, \*\**P* < 0.01, \*\*\**P* < 0.001, \*\*\*\**P* < 0.0001 and ns indicates non-significance by unpaired Student’s *t* test (between two samples), and one-way ANOVA with post hoc Tukey’s test (multiple comparisons).

### TFSA NPs increase mitochondria turnover and improve mitochondrial morphology and functions in INS-1 β-cells under lipotoxic condition

The proper maintenance of mitochondrial turnover is needed to sustain normal mitochondrial functions. To determine whether increasing autophagic function with treatment of TFSA NPs improves the turnover of mitochondria, we examined mitophagy by transfecting INS-1 β-cells with a mCherry-GFP-FIS1 plasmid that reflects the extent of mitochondria accumulation inside lysosomes by observing their colocalization (37). In addition, when mitochondria are recruited into sufficiently acidified lysosomes, the low pH quenches the green fluorescent protein (GFP) signal without affecting mCherry fluorescence (37). In BSA treated control INS-1 β-cells, the mitophagy reporter shows a basal red and green fluorescence, indicating normal turnover of mitochondria. Under lipotoxic condition with palmitate treatment, the increase in the green fluorescence intensity signifies the poorly acidified lysosomes and the increased accumulation of mitochondrial proteins (**Figure 5A-B**). Together with the blue fluorescence from Lysosensor blue indicative of lysosomes, the white puncta formed by the overlap of red, green, and blue fluorescence indicate co-localization within the lysosomes (**Figure 5A-B**). Addition of TFSA NPs lowers the lysosomal pH, making them more acidic and thereby quenching the GFP fluorescence, leading to the appearance of purple puncta when colocalizing with lysosomes (**Figure 5A-B**). This result indicates that treatment with TFSA NPs promotes mitophagy and turnover of damaged mitochondria.

**Figure 5.**
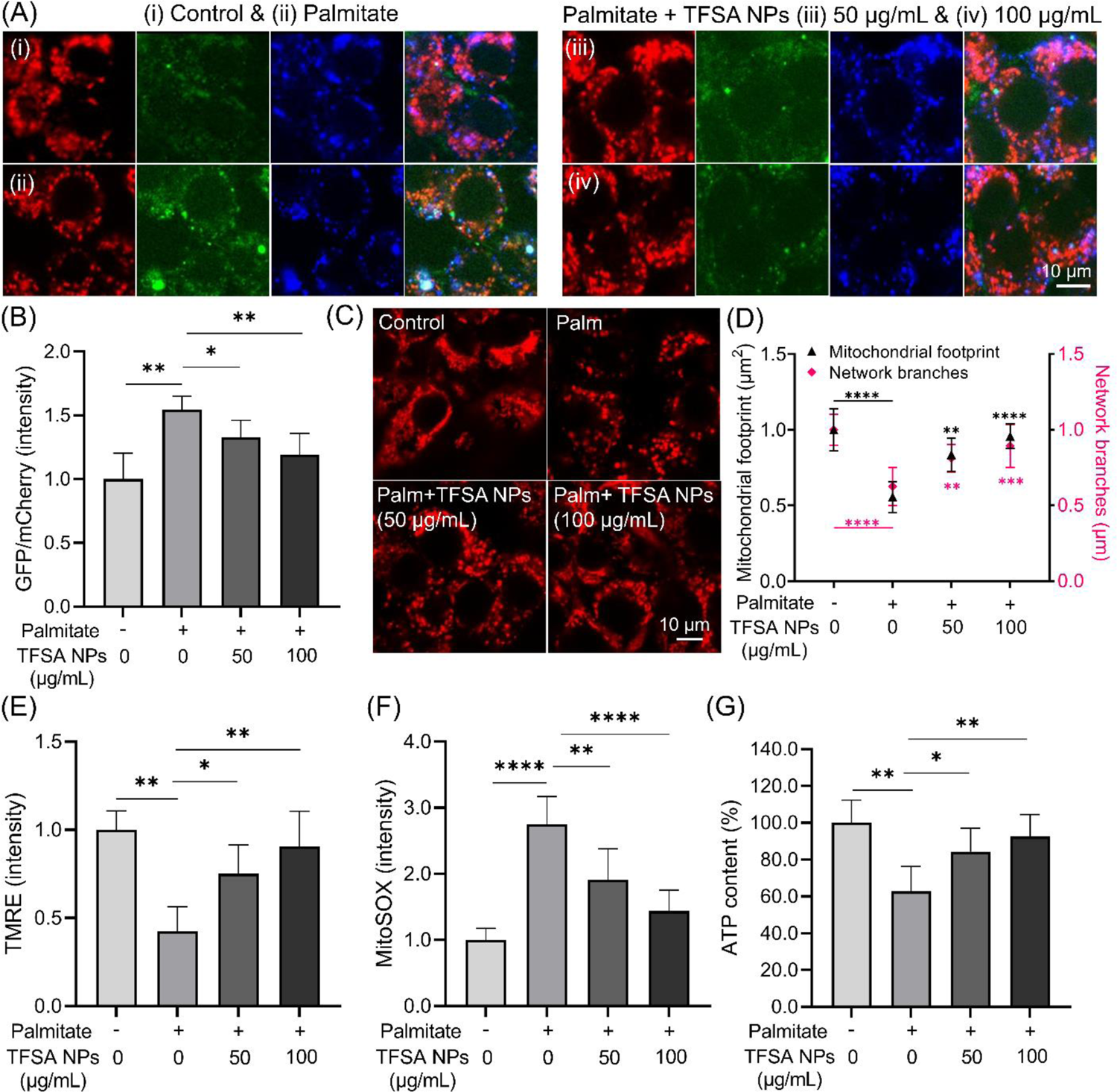
TFSA NPs improve mitochondrial turnover and functions in INS-1 β-cells under lipotoxic condition with palmitate treatment. (A-B) Confocal microscopy images of INS-1 β-cells transfected with mCherry-GFP-FIS1 mitophagy reporter plasmid followed by treatment with (i) BSA, (ii) palmitate, and palmitate with (iii) 50 µg/mL and (iv) 100 µg/mL of TFSA NPs. (C-D) Confocal images of INS-1 β-cells stained with MitoTracker Deep Red and quantification of mitochondrial footprint and network branches using MiNA analysis for samples treated with BSA, palmitate (palm), and palmitate with 50 µg/mL and 100 µg/mL of TFSA NPs. (E-G) Measurement of (E) mitochondrial membrane potential, (F) reactive oxygen species (ROS) generation, and (G) cellular ATP content under respective treatment conditions in INS-1 β-cells. Data presented are relative to control without palmitate and TFSA NPs. Data are means ± SD of N=4 independent experiments. \**P* < 0.05, \*\**P* < 0.01, \*\*\**P* < 0.001 and \*\*\*\**P* < 0.0001 by one-way ANOVA with post hoc Tukey’s test (multiple comparisons).

To determine whether TFSA NPs restore mitochondrial functions, we first examined the quality of mitochondria under palmitate treatment without and with TFSA NPs addition using MitoTracker Deep Red. Palmitate treatment induces more mitochondrial fragmentation as illustrated from the decrease in mitochondrial footprint and network branches (**Figure 5C-D**), which are associated with the impairment of mitochondrial functions (38, 39). TFSA NPs promote the presence of more elongated mitochondria with tubular mitochondrial network (**Figure 5C-D**), indicating an improvement in mitochondrial functions. We further probed for mitochondrial functions and demonstrated that TFSA NPs restore mitochondrial membrane potential (MMP) as shown by an increase in TMRE intensity (**Figure 5E**), reduce ROS generation (**Figure 5F**), and increase cellular ATP content (**Figure 5G**) as compared to palmitate treated INS-1 β-cells without TFSA NPs treatment.

### TFSA NPs promote glucose-stimulated insulin secretion in palmitate-treated INS-1 β-cells and increase glucose clearance in HFD mouse model of T2D

To determine if TFSA NPs modulate pancreatic β-cells functions, we first investigated if TFSA NPs affect glucose-stimulated insulin release from the INS-1 β-cells. Palmitate treatment reduces the amount of insulin secretory granules upon glucose stimulation, and TFSA NPs reverse this process to induce the production of more intracellular insulin vesicles in INS-1 β-cells (**Figure 6A-B**). We further confirmed that palmitate treatment reduced insulin secretion in INS-1 β-cells upon glucose stimulation, and that the addition of TFSA NPs restores and promotes insulin secretion (**Figure 6C**). Finally, we investigated the effect of TFSA NPs in a HFD mouse model of T2D. First, TFSA NPs treatment did not change the body weight (**Figure S6A**) and food intake (**Figure S6B-C**) in both control and HFD mice. We then showed that mice on 12 weeks of HFD had high fasting plasma insulin and blood glucose levels (**Figure 6D-E**), which are associated with insulin resistance observed in T2D patients. Importantly, intravenous administration of TFSA NPs lowers blood insulin and glucose levels (**Figure 6D-E**) and significantly promotes glucose clearance (**Figure 6F**), indicative of an overall improvement in glucose utilization by the body. To account for differences in baseline fasting glucose levels measured across the treatment groups, we calculated the area under the glucose tolerance response curve, and TFSA NPs significantly decrease total glucose excursion compared to HFD mice without TFSA NPs treatment (**Figure 6G**). It is important to note that there is no significant effect of TFSA NPs on the control mice. Together, our data suggest that TFSA NPs enhance lysosomal acidification in pancreatic β-cells under lipotoxic condition to improve lysosomal and autophagic functions and increase turnover of impaired mitochondria via mitophagy, thus promoting the presence of functional mitochondria. Improved mitochondrial function subsequently produces more ATP and increases insulin secretion under glucose stimulation in pancreatic β-cells to promote the utilization of the excessive glucose that are present under T2D condition.

**Figure 6.**
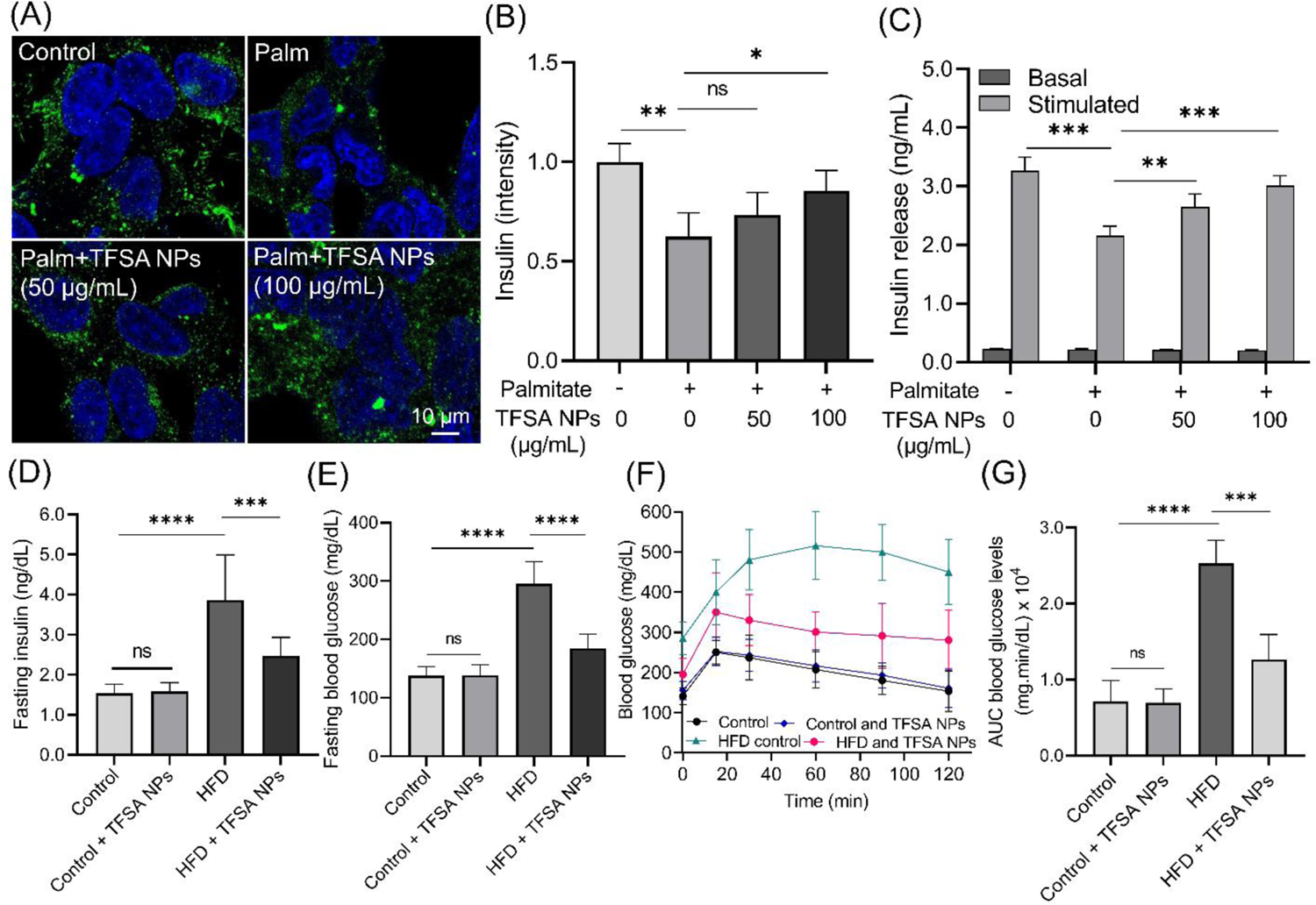
TFSA NPs promote insulin secretion in palmitate treated INS-1 β-cells and improve metabolic function in HFD mouse model of T2D. (A-B) Immunofluorescence and quantification demonstrating a decrease in intracellular insulin granules in INS-1 β-cells under lipotoxic condition with palmitate (palm) treatment and TFSA NPs increase the presence of these intracellular insulin granules. Data presented are relative to control without palmitate and TFSA NPs. (C) Insulin secretion is decreased in palmitate treated INS-1 β-cells and treatment with TFSA NPs increases glucose-stimulated insulin secretion. There is no statistical difference in basal insulin secretion levels among all treatment conditions. (D) Measurement of fasting blood insulin levels in control mice without and with treatment of TFSA NPs, and HFD mice without and with treatment of TFSA NPs. (E) Measurement of fasting blood glucose levels in control mice without and with treatment of TFSA NPs, and HFD mice without and with treatment of TFSA NPs. (F-G) Assessment of glucose tolerance by measuring (F) glucose tolerance response and (G) total glucose excursion in control mice without and with treatment of TFSA NPs, and HFD mice without and with treatment of TFSA NPs. Data are means ± SD of N=4 independent cell experiments, and N=8 mice per treatment group. \**P* < 0.05, \*\**P* < 0.01, \*\*\**P* < 0.001, \*\*\*\**P* < 0.0001 and ns indicates non-significance by one-way ANOVA with post hoc Tukey’s test (multiple comparisons).

Our results illustrate for the first time the reduction of lysosomal V-ATPase subunits that contributes to defective lysosomal acidification in INS-1 β-cells under lipotoxic conditions. This is supported by our recent studies demonstrating that restoration of lysosomal acidification improves autophagic and mitochondrial functions in INS-1 β-cells (16, 20) and in a HFD induced mouse model of T2D and NAFLD (40, 41), although the level of lysosomal subunits was not investigated in these studies. Complementary to our previous work illustrating the enhancement of lysosomal function in hepatocytes to stimulate insulin receptor signaling and mediate insulin resistance (40), our current study shows that promoting lysosomal acidification and function by TFSA NPs in β-cells increases insulin secretion to mediate T2D. However, it is worth noting that there might be other contributing factors to the restorative effect of TFSA NPs in promoting glucose utilization in the HFD mice such as promoting insulin receptor signaling to mediate insulin resistance in other organs due to the nature of this characterization *in vivo*. For example, insulin injection has been shown to upregulate lysosomal V-ATPase subunits (ATPV1A, ATPV1B2, ATPV1F, ATPV1E1, ATPV0A1) in the livers of Sprague-Dawley T2D rats (42). In addition, inhibition of lysosomal V-ATPase function or acidification by bafilomycin A1 impairs insulin receptor recycling and insulin signaling (42), which are implicated in causing insulin resistance (43, 44). Hence, re-acidification of impaired lysosomes could potentially increase insulin receptor recycling and signaling, thereby reducing insulin resistance in T2D. Furthermore, lysosomal dysfunction has been suggested as a trigger for inflammation in obesity (45, 46), which could induce further undesired complications.

Studies conducted in non-obese diabetic (NOD) mice also support the role of lysosomal acidification dysfunction in diabetes where there is a reduction of lysosomal subunit ATP6V1A in the mouse islets (47, 48). In fact, deletion of ATP6AP2 leads to β-cell lysosomal acidification dysfunction, autophagic function impairment, and defective insulin granule turnover and secretion (49). While it has been shown in T2D, it is unclear whether improving β-cell lysosome function, such as promoting its acidification, can prevent type 1 diabetes (T1D) development. It is also suggested that reduced insulin granule turnover will result in intracellular degradation of insulin, thus decreasing their secretion (49). This finding is consistent with another study which shows that palmitate treatment of INS-1 β-cells increases lysosomal localization and degradation of nascent insulin granules resulting in decreased insulin release from cells (50). We note that it is possible that lysosomal re-acidification in palmitate treated cells can also exert direct effects on the degradation of nascent insulin granules, which counteracts its ability to restore insulin secretion due to the restoration of autophagic and mitochondrial functions. However, our results show an increase in insulin release from INS-1 β-cells treated with palmitate and TFSA NPs, suggesting that re-acidification of impaired lysosomes to restore autophagic and mitochondrial functions plays a dominant role in the recovery of insulin secretion function of pancreatic β-cells.

In pancreatic β-cells, maintenance of mitochondrial oxidative phosphorylation increases cellular ATP content which induces depolarization of the plasma membrane via closure of ATP-sensitive K^+^ channels. Subsequent opening of voltage-dependent Ca^2+^ channels leads to an increase in intracellular Ca^2+^, which triggers the release of insulin from insulin-containing secretory granules (51). Importantly, increased cellular ROS level can lead to damages in mitochondrial complexes by oxidative stress and reduce the capability of mitochondria in producing ATP (52, 53), contributing decreased glucose-stimulated insulin secretion (54). Our current work, together with previous studies, demonstrate that palmitate disrupted mitochondrial network, reduced MMP, impaired mitochondrial function, increased ROS generation, and significantly decreased cellular ATP content (14, 55–57). Re-acidification of impaired lysosomes with TFSA NPs increased turnover of defective mitochondria, reduced ROS generation, restored cellular ATP content, and promoted glucose-stimulated insulin secretion by β-cells, which is consistent with our previous observation (16).

NPs are an increasingly important and useful tool for targeting and restoring lysosomal acidification (20–22, 58, 59). Here, we present a new type of TFSA NPs containing a single TFSA carboxylic acid as its constituent, which offers several advantages over the other NPs composed of a mixture of carboxylic acids. Using a single carboxylic acid enables finer control over composition and the molecular weight of the resulting polyester, hence generating polymers with a narrower distribution of molecular weights and resulting in more consistent and predictable material properties. In addition, acquisition of single carboxylic acid can be more cost-effective which presents a more practical choice for future large-scale productions and applications. Besides polymer based therapeutics, small molecules are being used to modulate lysosomal pH (21), although their effects have not yet been tested in obesity related metabolic disorders, such as T2D and NAFLD. To identify more effective modulators of lysosomal pH, drug screening studies can be conducted, such as using artificial lysosomes loaded with trypsin (60, 61).

## Conclusion

In conclusion, we provide the first evidence, from both bioinformatics and experimental analyses, that downregulation of lysosomal V-ATPase subunits occurs in both human T2D islets and palmitate induced pancreatic INS-1 β-cell model of T2D. This finding directly strengthens the significance of lysosome acidification impairment in disease pathogenesis and supports the continued development of acidifying therapeutics. To that end, we describe new biodegradable polymers based on TFSA and 1,4-butanediol and fabricate lysosome acidifying NPs (TFSA NPs). TFSA NPs treatment rescues decreased lysosomal acidity due to reduced subunit expression and restores autophagic and mitochondrial functions, leading to increased insulin secretion from pancreatic β-cells. TFSA NPs represent a new class of lysosome acidifying therapeutics applicable to T2D and obesity induced metabolic diseases in general, including Alzheimer’s disease which has been proposed to be type 3 diabetes.

## Materials and Methods

### Polyesters synthesis and characterization

The series of PBFSU polyester is synthesized via the following procedures. All reaction flasks were oven dried overnight prior to use. Di-acid monomers TFSA (Matrix Scientific, Cat# 006154) and SA (Sigma Aldrich, Cat# 398055) were added at varying ratios (TFSA:SA at 0:100, 25:75, 50:50, 75:25 or 100:0) in a round bottom flask. The linker 1,4-butanediol (Sigma Aldrich, Cat# 240559) was added at a 10 mol % excess, together with metal catalyst titanium(IV) isopropoxide (TIPT) (Sigma Aldrich, Cat# 377996), and distilled azeotropically at 120 °C for 16 h. Subsequently, a vacuum was slowly applied to prevent excessive foaming and minimize oligomer sublimation and allow for further condensation to form higher molecular weight polymer chains. The temperature was further increased to 130–140°C for at least 12 h. Finally, the viscous liquid or solid was precipitated in cold diethyl ether and dried under high vacuum for storage and further use. No unexpected or unusually high safety hazards were encountered. ^1^H-NMR, ^13^C-NMR and ^19^F-NMR spectra were recorded on a Varian INOVA 500 MHz spectrometer where chloroform (CDCl_3_) was used as the solvent. GPC was used to determine polymer molecular weights using tetrahydrofuran (THF) as the eluent and running against a narrow polystyrene standard curve (Aligent, PS-1, Standard B) at a flow rate of 1.0 mL/min through two Jordi columns (Jordi Gel DVB 10^5^ Å and Jordi Gel DVB 10^4^ Å, 7.8 × 300 mm) at 25 °C in series with a refractive index detector.

### Differential scanning calorimetry (DSC) and thermogravimetric analysis (TGA)

The thermal transitions were recorded with DSC on a Q200 thermal analyzer (TA Instruments) with a standard heating−cooling−heating mode. The heating rate is 10 °C/min and cooling rate is 5 °C/min. The thermal decomposition behavior was recorded with TGA (Q500, TA Instruments). The samples were heated from room temperature to 600 °C at 20 °C/min under N_2_ atmosphere.

### NPs synthesis and characterization

The formation of the NPs from polyesters was performed through a nanoprecipitation method (62). Briefly, the polyesters (PBSU, 25% PBFSU, 50% PBFSU, 75% PBFSU and 100% PBFSU) are dissolved in acetonitrile (Sigma Aldrich, Cat # 271004) and filtered through a 0.2 µm syringe filter (Millipore, Cat# Z741696) to remove particulates. Sodium dodecyl sulfate (SDS) (Sigma Aldrich, Cat# L3771) is dissolved in MilliQ water and stirred at high speed at 1700 rpm. The polyester solution is then added drop wise into the fast-stirring aqueous solution. Immediately after adding all the polyester solution, the emulsion is placed into a SnakeSkin dialysis tubing (MWCO 10 kDa) (Thermofisher Scientific, Cat# 68100) and dialyzed against MilliQ water for 24 h. For dynamic light scattering (DLS) measurements, 200 µL of the solution is diluted in 2.8 mL of MilliQ water, and the diameter and zeta potential are obtained from the Brookhaven 90 plus NP sizer DLS instrument.

### Scanning electron microscopy (SEM)

NPs were diluted 50 times in MilliQ water and 10 µL aliquots were plated on silicon wafers and allowed to air dry overnight. The wafers were then affixed to aluminum stubs with copper tape and sputter coated with 5 nm Au/Pd. These samples were then imaged using a Supra 55VP field emission scanning electron microscope (ZEISS) with an accelerating voltage of 2 kV and working distance of 5.5 cm.

### NPs degradation assays

NPs (10 mg/mL) were diluted in 20 mM phosphate saline buffer (PBS) adjusted to pH 7.4 or 6.0. The pH changes due to NPs degradation is measured at different time intervals using a pH meter. At specific time points, the solutions containing NPs were centrifuged and the pellets were dried in N_2_ atmosphere overnight. The samples were then dissolved in THF and filtered with a 0.22 µm syringe filter. GPC was used to determine polymer molecular weights of the NPs using THF as the eluent following the same method as described above.

### Cell culture and MTS assay

INS-1 832/13 rat insulinoma β-cells (Millipore, Cat# SCC207) were cultured in RPMI media supplemented with 10% fetal bovine serum (FBS), 1mM glutamine, 50 µM β-mercaptoethanol, 50 units/mL penicillin, and 50 g/mL streptomycin. The cytotoxicity of the different types of PBFSU NPs were evaluated using an MTS Assay Kit (Abcam, Cat# ab197010). INS-1 β-cells were cultured in a 96-well plate at 15000 cells/well for 24 h and subsequently treated with the respective doses of the different types of PBFSU NPs for 24 hours. The cell viability was quantified relative to the no treatment control.

### Data mining of microarray transcriptome dataset

Microarray dataset GSE25724 containing mRNA expression of human T2D (6 diseased samples) and non-diabetic (7 healthy control samples) islets (33) was obtained from the Gene Expression Omnibus (GEO) database (63). Data mining was conducted by analyzing the GSE25724 dataset using the GEO2R tool and the limma package in the Rstudio (64, 65). The mRNA expression from 6 T2D samples were compared against the 7 non-diabetic samples, and the DEGs were identified using the parameters of adjusted *P*<0.05 (*P*-values corrected for multiple testing using false discovery rate (FDR) method) and log_2_FC≥|1|. To visualize the spread, significance, and difference in expression levels of the DEGs, volcano plots and heatmaps were generated using the ggplot2 and pheatmap packages in Rstudio, respectively. The pathway enrichment analysis was conducted using the Database for Annotation, Visualization and Integrated Discovery (DAVID) (66) which provides functional annotations of the DEGs based on KEGG and GOBP pathways.

### Palmitate:BSA preparation

Palmitic acid (Sigma Aldrich, Cat# P5585) was first dissolved in dimethyl sulfoxide (DMSO) (Millipore, Cat# D8418). 1 mL of the DMSO solution was then dissolved at 45 °C in 100 mL of RPMI media containing 6.7g of fatty acid-free BSA (Millipore, Cat# A7030) to make a 4 mM (10X) stock of palmitate:BSA complex. For BSA control, a 10X stock of 100 mL of RPMI media containing 6.7g of BSA and 1 mL DMSO (no palmitic acid) was made. For cell treatments, the 10X stocks were diluted in RPMI media with FBS to form 1X treatment media. The pH of the treatment media was then adjusted to 7.4 followed by sterile filtration before addition to the INS-1 β-cells for 24 h.

### Lysosomal pH quantification and cathepsin L activity assay

For lysosomal pH imaging, INS-1 β-cells under respective treatments were stained with 1 µM LysoSensor™ Yellow/Blue DND-160 (Thermofisher scientific, Cat# L7545) for 5 min followed by imaging using confocal microscope (ZEISS) at 360 nm excitation and acquiring images at 440 nm (blue channel) and 540 nm (yellow channel). The ratio between yellow and blue intensity was calculated and pH quantification was achieved by benchmarking against a standard curve with known pH values. The standard curve was established by matching LysoSensor fluorescence ratio to known pH using 2-(N-morpholino) ethanesulfonic acid buffer of varying pH. For the cathepsin L activity assay, INS-1 β-cells treated with respective conditions were stained with 10 μg/mL Magic red cathepsin L (MR-cathepsin L, Immunochemistry Technologies, Cat# 941) for 1 h. The cells were then washed three times with PBS and the fluorescence was measured with a Synergy H1 model of hybrid multi-mode plate-reader with excitation/emission 531/629 nm. The fluorescence was normalized to cell count for each condition.

### Mitochondrial morphology and turnover analysis

INS-1 β-cells treated with respective conditions were stained with 250 nM MitoTracker Deep Red (Thermofisher scientific, Cat# M22426) for 30 min, before imaging using confocal microscopy at excitation/emission 640/665 nm. For mitochondrial morphology analysis, acquired confocal images of mitochondria were preprocessed through ImageJ using the unsharp mask filter to sharpen the image and applying a median filter to limit background signals (67, 68). Image was then segmented into smaller tiles each containing a single cell to calculate individual mitochondrial morphology. Each tiled image was then processed through the Mitochondrial Network Analysis Image J macro (MiNA) (69), where they were converted into a binary image and skeletonized. An analysis of the skeleton will be recorded, followed by generation of a tabularized output. Quantifiable measurements of mitochondrial morphology, including mitochondrial footprint (µm^2^) and network branch length (µm) were then plotted. Parameters for processing were kept consistent across images acquired from different treatment conditions. To quantify mitochondria turnover under different conditions, INS-1 β-cells were first transfected with mCherry-GFP-FIS1 (University of Dundee) for 24 h to express the mitophagy reporter followed by applying the different treatment conditions for another 24 h. The LysoSensor™ Blue DND-167 dye (Thermofisher scientific, Cat# L7533) was added for 15 min before imaging of the cells. Confocal microscope was used to acquire the fluorescent images at blue (excitation/emission 405/460 nm), green (excitation/emission 490/510 nm) and red (excitation/emission 590/610 nm) channels.

### Mitochondrial functional assays

INS-1 β-cells were cultured in 96-well plate and treated with respective conditions. Mitochondrial membrane potential was analyzed with TMRE-Mitochondrial Membrane Potential Assay Kit (Abcam, Cat# ab113852). Mitochondrial superoxide release was measured with Mitochondrial Superoxide Assay Kit (Abcam, Cat# ab219943), according to manufacturer’s instructions. Measurements were acquired using Synergy H1 plate-reader with excitation/emission 549/575 nm for TMRE and excitation/emission 510/610 nm for superoxide signals.

### Measurement of cellular ATP content

For cellular ATP content determination, INS-1 β-cells were cultured in a 6-well plate and treated with respective conditions for 24 h. Subsequently, the cells were washed twice with cold PBS and resuspended in the assay buffer provided for homogenization according to the manufacturer’s instructions (Abcam, Cat# ab83355). The supernatant of the cell lysates was incubated with ATP reaction mix for 30 min followed by fluorescence measurements using Synergy H1 plate-reader with excitation/emission 535/587 nm.

### Western blotting

The INS-1 β-cells treated with respective conditions were first washed with PBS, followed by cell lysis using RIPA buffer (Thermofisher Scientific, Cat# 89900) with 2% Triton-X-100 and protease and phosphatase inhibitors (Thermofisher Scientific, Cat# 78440). Lysates were kept on ice for 15–30 min followed by centrifugation for 10 min at 13,500 × g at 4 °C. Pierce™ BCA Protein Assay Kit (Thermofisher Scientific, Cat# 23225) was used to determine total protein concentrations for each sample. Samples were then mixed with 4X Laemmli Sample Buffer (Bio-Rad, Cat# 1610747) and boiled at 95 °C for 5 min before loading and running in 4-15% Mini-PROTEAN TGX Precast Protein Gels (Bio-Rad, Cat# 4561083) with Precision Plus Protein Dual Color Standards (Bio-Rad, Cat# 1610374). The proteins were transferred to polyvinylidene fluoride (PVDF) membrane (Millipore, Cat# ISEQ00010), and blocked with blocking buffer (Bio-Rad, Cat# 1706404). The membrane is then incubated with a series of primary and secondary antibodies. Primary antibodies used include: ATP6V1B2 (Proteintech, Cat# 15097-1-AP, RRID: AB_2243297, 1:1000), ATP6AP2 (Proteintech, Cat# 10926-1-AP, RRID: AB_2062201, 1:1000), ATP6V1A (Proteintech, Cat# 17115-1-AP, RRID: AB_2290195, 1:1000), LC3A/B (Cell Signaling, Cat# 12741; RRID: AB_2617131, 1:1000), SQSTM1/p62 (Cell Signaling, Cat# 5114; RRID: AB_10624872, 1:1000), and GAPDH (Cell Signaling, Cat# 2118; RRID: AB_561053, 1:1000). Secondary antibodies used include anti-rabbit IgG, HRP-linked antibody (Cell Signaling, Cat# 7074; RRID: AB_2099233, 1:3000) and anti-mouse IgG, HRP-linked antibody (Cell Signaling, Cat# 7076; RRID: AB_330924; 1:3000). Densitometry was performed using ImageJ and protein expression levels were normalized to GAPDH loading control.

### Insulin secretion assays (immunostaining and ELISA)

Prior to stimulating insulin secretion by glucose, the INS-1 β-cells were treated with respective conditions and undergone a 2 h culture in RPMI medium supplemented with 2 mM glucose and devoid of serum. Subsequently, the cells were rinsed and subjected to a 30 min preincubation in cell culture medium containing 2 mM glucose. Next, the cells were incubated for 1 h in cell culture media containing either 2 (low) or 16 (high) mM of glucose. To perform immunostaining of insulin, INS-1 β-cells were first fixed in 4% paraformaldehyde (PFA) in PBS solution (VWR, Cat# ALFAJ61899.AK) for 30 min, before washing twice with PBS. The fixed cells are then washed with 0.1% Triton X-100 (Sigma Aldrich, Cat# X100) in PBS for 3 times with 5 min each. The fixed cells are blocked with 3% normal goat serum (NGS) (Thermofisher Scientific, Cat# 31872) for 1 h at room temperature. The fixed cells were then incubated with primary antibodies against insulin (Cell Signaling, Cat# 3014, AB_2126503, 1:500) in 1% NGS solution overnight at 4 °C. The fixed cells were then washed with 0.1% Triton X-100 in PBS solution for 3 times with 5 min each and incubated with Goat anti-rabbit 488 nm (Thermofisher Scientific Cat# A-11008, RRID: AB_143165, 1:1000) fluorescent secondary antibody for 1 h at room temperature followed by washing with PBS. The fixed cells were then stained with DAPI (Thermofisher Scientific, Cat# 9542) for 10 min and washed with PBS. Fluorescent images were acquired using confocal microscope (ZEISS) at blue (excitation/emission 405/460 nm) and green (excitation/emission 490/520) channels. To quantify insulin secretion, enzyme linked immunosorbent assay (ELISA) assay against insulin was conducted. The cell culture media from the respective samples were collected for insulin measurement. The analysis of insulin secretion from INS-1 β-cells was conducted using the Insulin Sandwich ELISA Kit (Proteintech, Cat# KE20008), following the instructions provided by the manufacturer. Measurements were acquired using Synergy H1 plate-reader with an absorbance of 450 nm.

### Maintenance of animals

We utilized a well-established *in vivo* model involving in-house bred C57BL/6J male mice fed with a HFD (Altromin, Cat# C1090-60) for a duration of 12 weeks as a model for T2D. As a control group, C57BL/6 J male mice were fed with a normal chow diet. The mice were housed in a temperature-controlled environment (25 °C) within virus-free facilities that maintain a 12 h light/dark cycle (7:30 a.m. on/7:30 p.m. off) and had unrestricted access to water. For each animal experiment, the mice on either chow or HFD were randomly assigned to receive injections with either saline or 300 mg/kg of TFSA NPs via the tail vein. Injections were done 3 times, once every other day, for a period of 5 days. We closely monitored the mouse body weights and food intake for any initial signs of toxicity during the injection period. All mice received proper treatment in accordance with the approved protocols A22058 and A21043 from Nanyang Technological University (NTU) Institutional Animal Care and Use Committee (IACUC). Glucose tolerance tests were conducted 24 h after the final NPs injection.

### Animal studies (fasting plasma insulin and glucose levels/glucose tolerance test)

Mice were fasted for 6 h prior to conducting all experiments. All mouse blood samples were collected from the tail using Microvette capillary tubes (Sarstedt Inc, Cat# NC9141704). Fasting plasma insulin levels were measured using the Mouse Insulin ELISA Kit (Proteintech, Cat# KE10089). Fasting blood glucose concentrations were measured using blood glucose strips and a blood glucometer. For glucose tolerance test, an intraperitoneal injection of glucose at 2 g/kg body weight was administered. The blood glucose concentrations were measured at baseline as well as at 15, 30, 60, 90, and 120 min after the glucose injection.

### Statistical analysis

Statistical analyses were performed using the GraphPad Prism software. Unpaired Student’s *t* test was used to validate statistical difference between two conditions. One-way ANOVA with post hoc Tukey’s test was used when more than two conditions were analyzed. *P*<0.05 was considered statistically significant. \**P*<0.05, \*\**P*<0.01, \*\*\**P*<0.001, \*\*\*\**P*<0.0001 and ns indicates non-significance.

## Competing interest

M.W.G., O.S.S., and J.Z. have jointly invented and hold a patent (Patent number: US10925975B2) registered with the United States Patent and Trademark Office. This patent focuses on the utilization of acidic nanoparticles as a therapeutic approach for diseases characterized by compromised lysosomal acidity. O.S.S. and M.W.G. are also co-founders of Enspire Bio/Capacity Bio, which are testing the application of these acidic nanoparticles. The remaining authors declare no additional competing interests.

## Funding

C.H.L. was supported by a Lee Kong Chian School of Medicine Dean’s Postdoctoral Fellowship (021207-00001) from Nanyang Technological University (NTU) Singapore and a Mistletoe Research Fellowship (022522-00001) from the Momental Foundation USA. J.Z. was supported by a Presidential Postdoctoral Fellowship (021229-00001) from NTU Singapore and a BU Nano Cross-disciplinary fellowship from the BU Nano Center at Boston University.

## Author contributions

C.H.L. and J.Z. designed the experiments and directed the project. J.Z., K.M.L., and M.W.G. with assistance from C.H.L. synthesized the polymers, engineered the TFSA nanoparticles and performed characterizations. C.H.L. and L.M.O. conducted the data mining and bioinformatics analysis of mRNA dataset. C.H.L., G.W.Z.L., E.N.S., and J.Z. carried out all cell-based assays and animal studies. C.H.L., K.M.L., J.I., and J.Z. performed data analysis. O.S.S. and M.W.G. provided critical comments and edited the manuscript. C.H.L. and J.Z. wrote the manuscript.

## Acknowledgements

The authors thank the funding sources for supporting this work.

---

**Figure S1.**
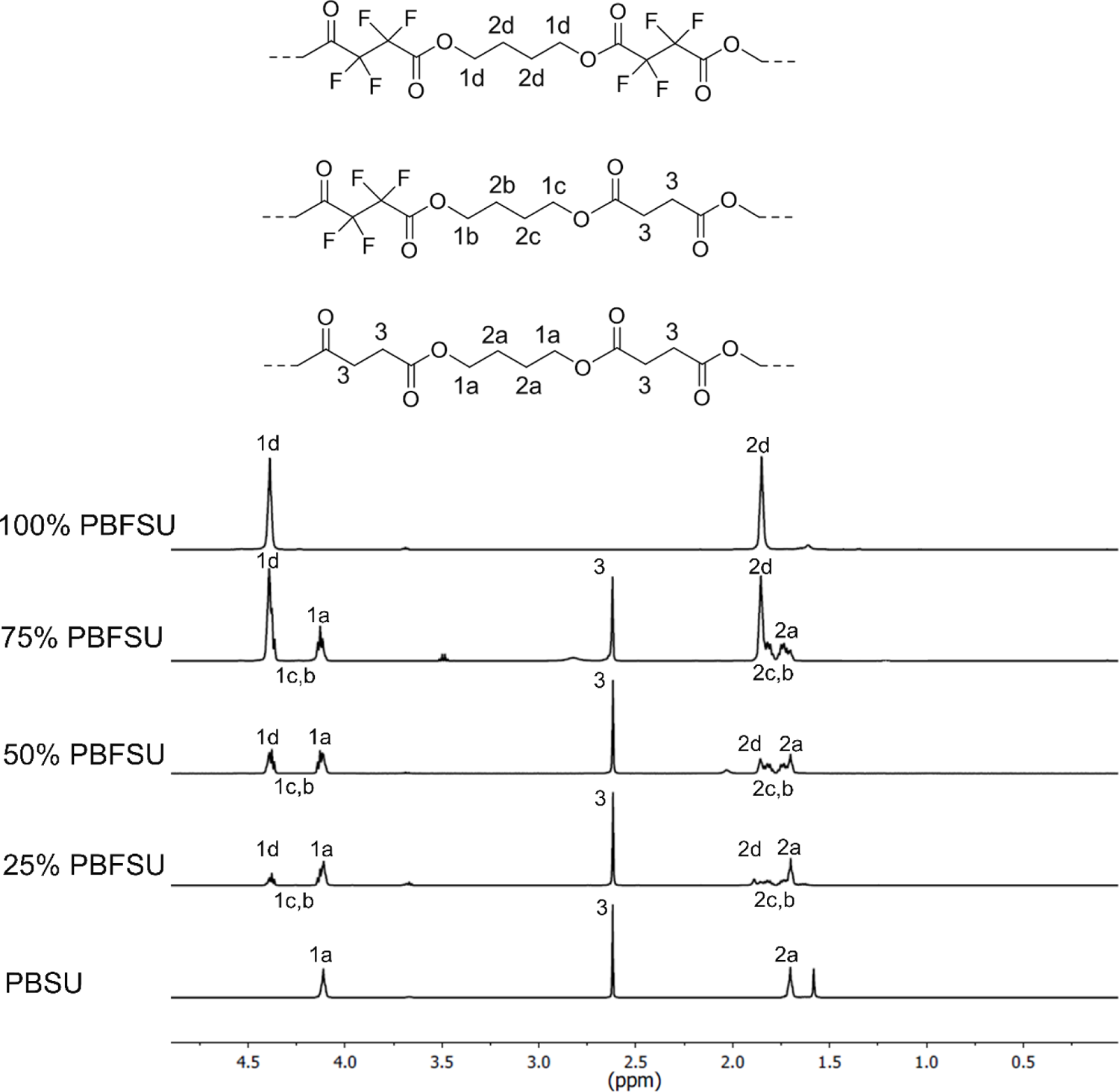
^1^H NMR spectra of PBFSU polyesters. ^1^H NMR [(500 MHz, CDCl_3_): [PBSU δ 1.58 (s, 1H), 1.70 (s, 2H), 2.62 (s, 2H), 4.11 (s, 2H)], [25% PBFSU 1.70 (m, 2H), 1.86 (m, 2H), 2.62 (s, 2H), 4.11 (m, 2H), 4.38 (m, 1H)], [50% PBFSU 1.70 (m, 2H), 1.86 (m, 2H), 2.62 (s, 2H), 4.11 (m, 2H), 4.38 (m, 2H)], [75% PBFSU 1.70 (m, 0.3H), 1.86 (m, 1H), 2.62 (s, 0.3H), 4.11 (m, 0.4H), 4.38 (m, 1H)], [100% PBFSU 1.86 (s, 1H), 4.39 (s, 1H)].

**Figure S2.**
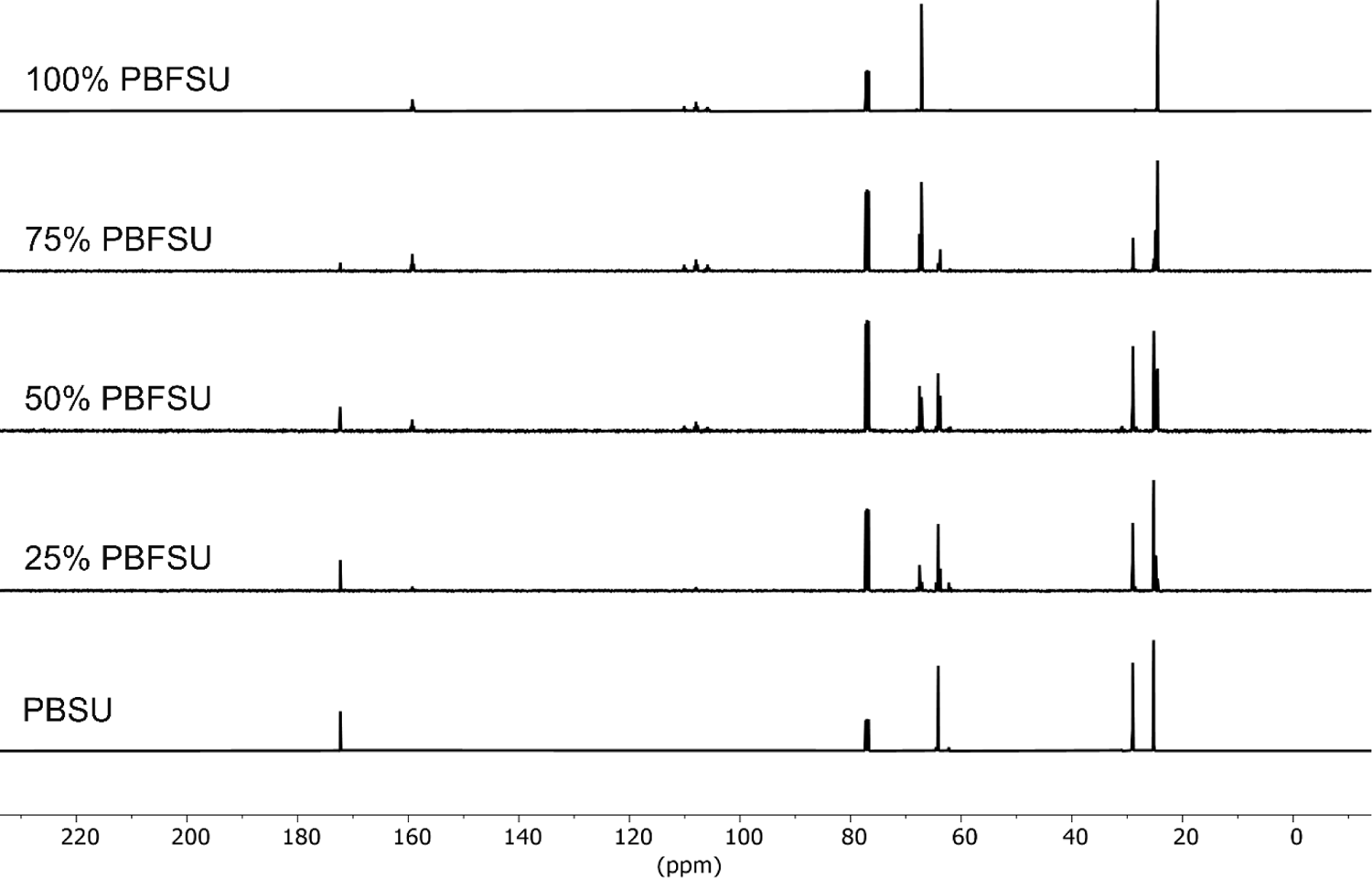
^13^C NMR spectra of PBFSU polyesters. ^13^C NMR [(500 MHz, CDCl_3_): [PBSU, 25.19, 28.99, 64.15, 77.28, 172.33], [25% PBFSU 25.18, 28.99, 67.52, 76.77, 107.76, 159.0, 172.28], [50% PBFSU 25.17, 28.95, 64.18, 67.52, 76.76, 107.98, 159.29, 172.09], [75% PBFSU 24.50, 28.92, 63.78, 67.17, 76.75, 107.76, 159.24, 172.31], [100% PBFSU 24.48, 67.17, 76.74, 107.97, 158.79].

**Figure S3.**
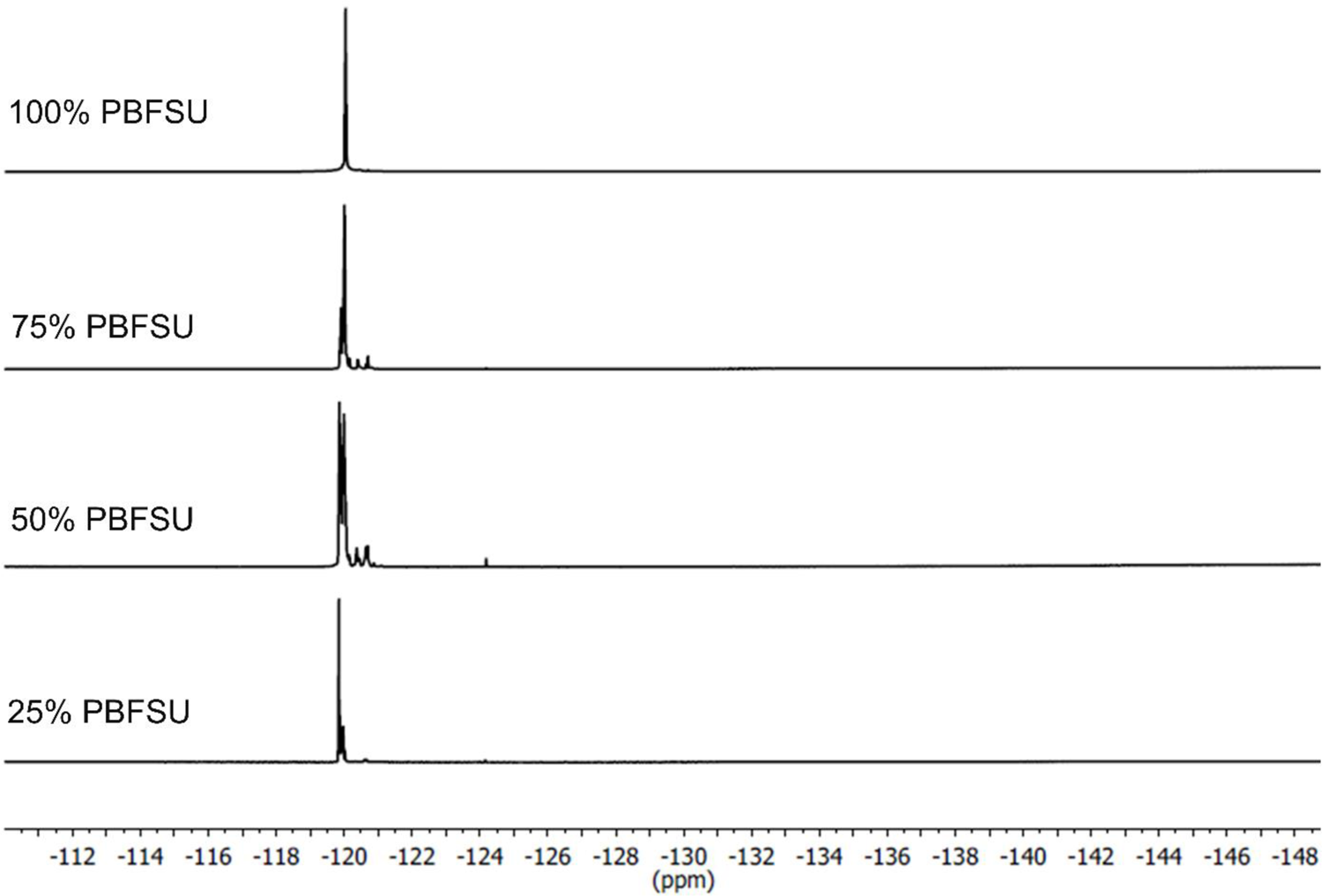
^19^F NMR spectra of PBFSU polyesters. ^19^F NMR [(500 MHz, CDCl_3_): [25% PBFSU-120.61, −120.02, −119.97, −119.85], [50% PBFSU −124.18, −121.40, −119.28], [75% PBFSU - 120.70, −120.40, −120.01], [100% PBFSU −120.04].

**Figure S4.**
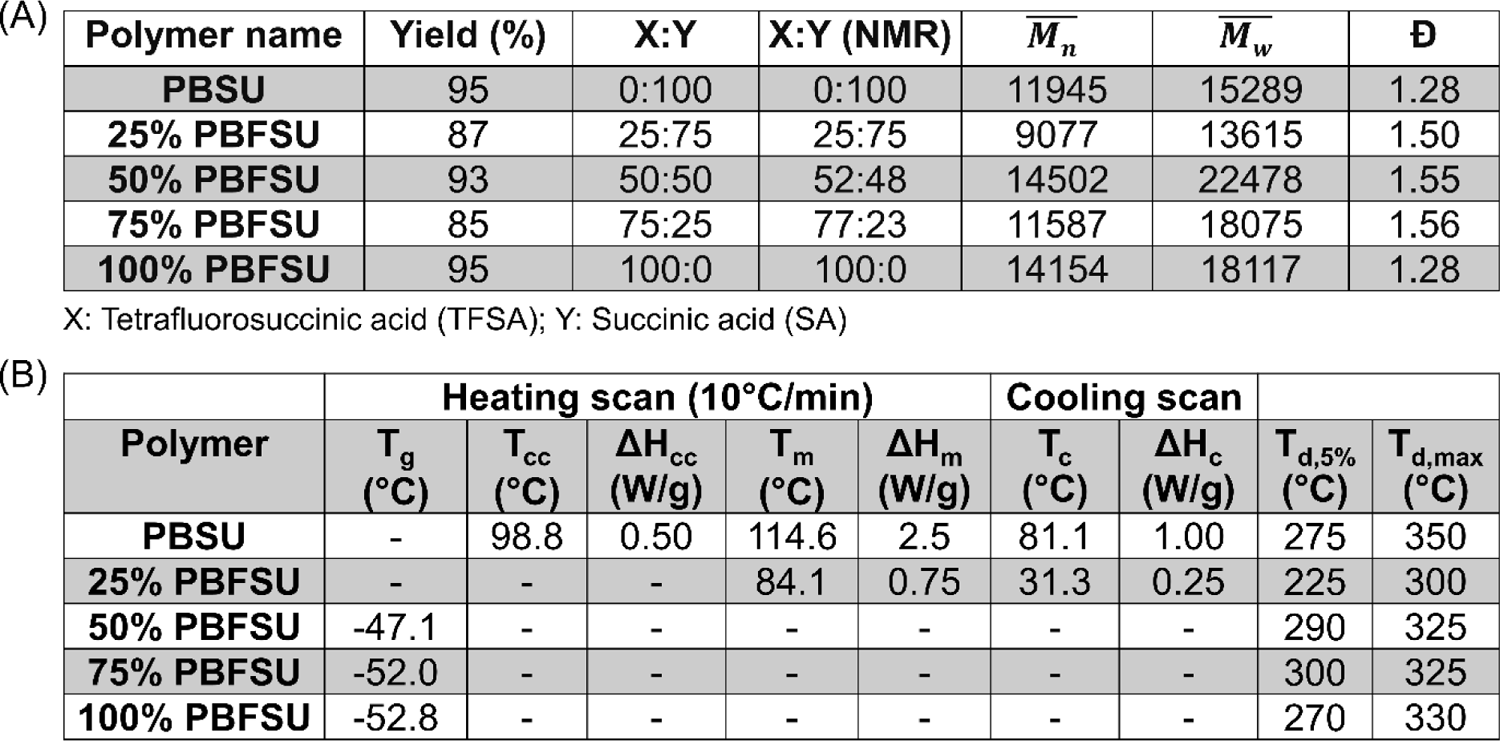
Characterization of the chemical and thermal properties of PBFSU polyesters. (A) Characterization of the percentage of the component acids using nuclear magnetic resonance (NMR), and number average molecular weights (*Mn*^-^), weight average molecular weights (*Mw*^-^) and polydispersity (Đ) of PBFSU polyesters using gel permeation chromatography (GPC). (B) Characterization of the thermal properties of PBFSU polyesters using differential scanning calorimetry (DSC) and thermogravimetric analysis (TGA). T_g_: transition temperature, T_cc_: cold crystallization temperature, ΔH_cc_: heat flow changes in cold crystallization, T_m_: melting temperature, ΔH_m_: heat flow changes in melting, T_c_: crystallization temperature, ΔH_c_: heat flow changes in crystallization, T_d,5%_: temperature at which polyester shows 5% decomposition, and T_d,max_: temperature at which the polyester shows maximum decomposition, and “–” indicates not applicable.

**Figure S5.**
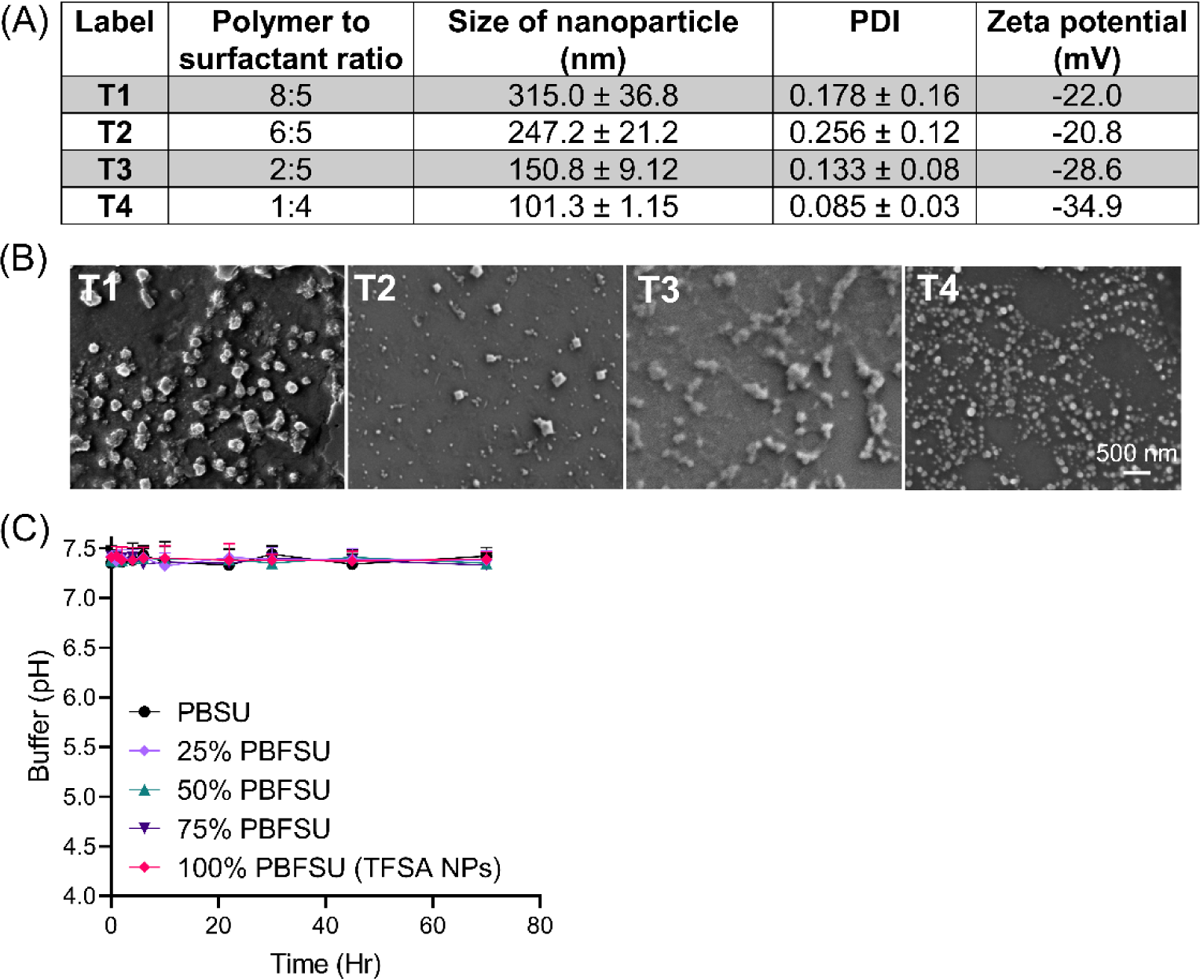
Optimization of the conditions for the formation of PBFSU based nanoparticles (NPs). (A) Optimization of the polymer to surfactant ratio for the formation of PBFSU based NPs.(B) Scanning electron microscopy (SEM) images of the PBFSU based NPs formed under different conditions in (A). The most optimized condition used to make the NPs is 1:4 polymer to surfactant ratio, reaction temperature at 25 °C and dialysis time of 24 h. (C) TFSA NPs (100% PBFSU), together with all other NPs, do not impart acidifying property under neutral condition of pH 7.4. Data are means ± SD of N=3 independent experiments.

**Figure S6.**
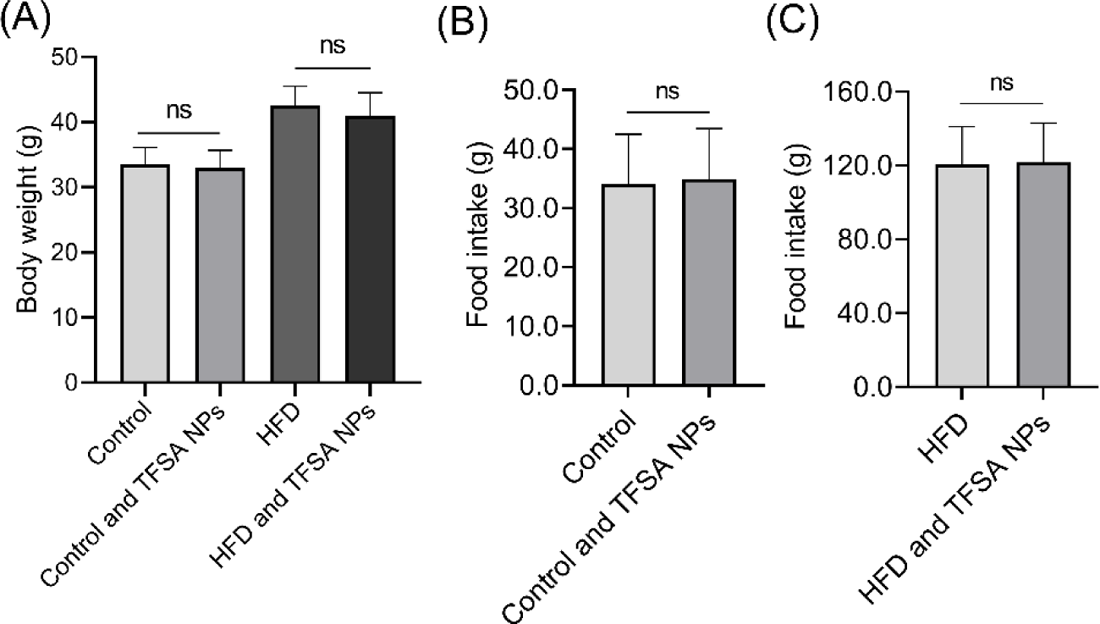
Characterization of body weight and food intake in control and HFD mice. (A) Body weight measurement of control mice without and with treatment of TFSA NPs, and HFD mice without and with treatment of TFSA NPs. (B) Food intake of control mice without and with treatment of TFSA NPs. (C) Food intake of HFD mice without and with treatment of TFSA NPs. Data are means ± SD of N=8 mice per treatment group. ns indicates non-significance by unpaired Student’s *t* test (between two samples), and one-way ANOVA with post hoc Tukey’s test (multiple comparisons).

## Notes

### Summary of Updates

Data mining of the T2D dataset was reanalyzed with adjusted p-value < 0.05. Three additional figure panels were added in the revised manuscript.

